# High affinity Na^+^ transport by wheat HKT1;5 is blocked by K^+^

**DOI:** 10.1101/280453

**Authors:** Bo Xu, Maria Hrmova, Matthew Gilliham

**Affiliations:** Australian Research Council Centre of Excellence in Plant Energy Biology, University of Adelaide, Waite Research Precinct, Glen Osmond, South Australia 5064, Australia; School of Agriculture, Food and Wine, and Waite Research Institute, University of Adelaide, Waite Research Precinct, Glen Osmond, South Australia 5064, Australia

**Keywords:** salinity, High-affinity K Transporters, membrane transport, KtrB, dual affinity, sodium exclusion

## Abstract

The wheat sodium transporters TmHKT1;5-A and TaHKT1;5-D are encoded by genes underlying major shoot Na^+^ exclusion loci *Nax2* and *Kna1* from *Triticum monococcum* (Tm) and *Triticum aestivum* (Ta), respectively. In contrast to HKT2 transporters that have been shown to exhibit high affinity K^+^-dependent Na^+^ transport, HKT1 proteins have, with one exception, only been shown to catalyse low affinity Na^+^ transport and no K^+^ transport. Here, using heterologous expression in *Xenopus laevis* oocytes we show that both TmHKT1;5-A and TaHKT1;5-D encode dual (high and low) affinity Na^+^-transporters with the high-affinity component being abolished when external K^+^ is in excess of external Na^+^. Based on 3-D structural modelling we propose that tighter binding of K^+^, compared to that of Na^+^ in the selectivity filter region by means of additional van der Waals forces, explains the K^+^ block at the molecular level. The low-affinity component for Na^+^ transport of TmHKT1;5-A had a lower K_m_ than that of TaHKT1;5-D and was less sensitive to external K^+^. We propose that these properties underpin the improvements in shoot Na^+^-exclusion and crop plant salt tolerance following the introgression of TmHKT1;5-A into diverse wheat backgrounds.

## Introduction

Crops suffer reduced growth and productivity under salinity stress. Salt, when it builds up to high concentrations in the growing medium (i.e. soil solution) imposes an osmotic limitation on water uptake, interferes with optimal nutrient homeostasis, and leads to the build up of leaf cellular sodium (Na^+^) concentrations, which causes an ionic toxicity that limits photosynthesis and carbon assimilation in plants (1).

A major genetic mechanism contributing towards salinity tolerance of most crops including the cereals rice (*Oryza sativa*) and wheat (*Triticum aestivum* and *Triticum monococcum*) is Na^+^ exclusion from leaves, with the High-affinity potassium (K^+^) transporter (HKT) protein family having a major role in this trait (2-6). This family of proteins is present in all plants so far sequenced; more broadly the members of the high affinity K^+^/Na^+^ transporting Ktr/TrK/HKT superfamily of proteins are present in bacteria (Ktr and TrK), fungi (TrK) and plants (HKT) (7,8).

These plant HKTs are divided into 2 clades: 1) class 1 HKT proteins (HKT1;x) are mostly Na^+-^selective transporters; 2) whereas class 2 HKTs (HKT2;y) mostly function as K^+^-Na^+^ symporters and so far have been only identified in cereal monocots (9,10). This definition has been challenged on occasion, such as for OsHKT2;4, where following its characterization in *Xenopus laevis* oocytes, there are conflicting reports on its permeability to Ca^2+^, Mg^2+^ and NH_4_^+^ in addition to K^+^ and Na^+^ (11-13). Furthermore, there are two reports of K^+^ permeability in the HKT1 clade. Firstly, *Arabidopsis* AtHKT1 could complement K^+^ transport deficient *E. coli*, although no K^+^ permeability could be found when *AtHKT1* was expressed in *X. laevis* oocytes in the same study (14). Secondly, *Eucalyptus camaldulensis* EcHKT1;1 and EcHKT1;2, which appear to be the exception for the HKT1 proteins characterized so far in that they transport both Na^+^ and K^+^ when expressed in *X. laevis* oocytes (15). Other HKT1 transporters have no reported K^+^ permeability (4,5,10,14,16-20).

Whilst K^+^ transport is not a common feature of HKT1 transporters, several have shown the property of K^+^-regulated Na^+^ transport (19,21-23). For instance, the presence of 10 mM external K^+^ ([K^+^]_ext_) reduced the inward Na^+^ current of *X. laevis* oocytes expressing *HKT1;5* from *T. aestivum* (*TaHKT1;5-D*) (19), and reduced outward Na^+^ currents generated by expression of *HKT1;2* from *Solanum lycopersicum* (*SlHKT1;2*) (21). Interestingly, [K^+^]_ext_ stimulated both inward and outward Na^+^ transport of *X. laevis* oocytes expressing two *HKT1;4* alleles from *Triticum turgidum* L. subsp. *Durum*, when assayed with solutions of very low ionic strength (23).

Ktr/TrK/HKT proteins are predicted to consist of 8 transmembrane α-helices that fold into four pseudo-tetramers around a central pore (6,16,18). The selectivity filter lining the pore of Ktr and TrK and class 2 HKT proteins is composed of four glycine residues. In class 1 HKTs the first glycine within the predicted selectivity filter is substituted with serine (16); the lack of K^+^ permeability in this clade has been linked to the presence of this glycine substitution (glycine for serine) (10,14,16-18); but the exact molecular mechanism underlying this block is to be determined.

Here, we further characterise HKT1;5 proteins underlying two major salt tolerance associated loci, *TmHKT1;5-A* (*Nax2*) and *TaHKT1;5-D* (*Kna1)* from *T. monococcum and T. aestivum* respectively. We find that both TmHKT1;5-A and TaHKT1;5-D encode dual affinity Na^+^-transporters and that their dual affinity Na^+^ transport can be blocked by raising the external K^+^ concentration. We further propose that the adjoining residues within the selectivity filter cause this block by the virtue of a higher number of van der Waals interactions with the K^+^ ion, than with Na^+^. Thus, the Na^+^ ion is capable of traversing the selectivity filter, while K^+^ causes a block. This is the first report that Na^+^ transport has two affinities in the HKT1 clade and demonstrates that dual affinity transport is a common property of both clades of the HKT family.

## Results

### K^+^ sensitivity of TmHTK1;5-A and TaHKT1;5-D

Previously, it was shown that inward Na^+^ transport of both TmHKT1;5-A and TaHKT1;5-D was inhibited by external K^+^ ([K^+^]_ext_) (4,19). Here, we further compared the K^+^-sensitivity of TmHKT1;5-A and TaHKT1;5-D (Fig. 1). An increase in [K^+^]_ext_ to 30 mM reduced the channel inward conductance (i.e. <100 mV) of TmHKT1;5-A by approximately 75% in a 1 mM Na^+^ ([Na^+^]_ext_) bath solution; similarly this amount of K^+^ inhibited that of TaHKT1;5-D up to 95 % (Figs. 1A, 1B and 2). However, such Na^+^-inhibition by K^+^ was gradually decreased by increasing [Na^+^]_ext_ (Figs. 1 and 2). For instance, 10 mM [K^+^]_ext_ suppressed the channel inward conductance of TmHKT1;5-A by 43.5% in 1 mM [Na^+^]_ext_ and by 16.5% in 10 mM [Na^+^]_ext_; it similarly reduced the TaHKT1;5-D inward conductance by 57% in 1 mM [Na^+^]_ext_ and by 23% in 10 mM [Na^+^]_ext_ (Figs. 1 and 2). When [Na^+^]_ext_ was increased to 30 mM, the channel inwared conductance of TmHKT1;5-A was insensitive to [K^+^]_ext_, whereas that of TaHKT1;5-D was inhibited (Fig. 2).

**Figure 1.**
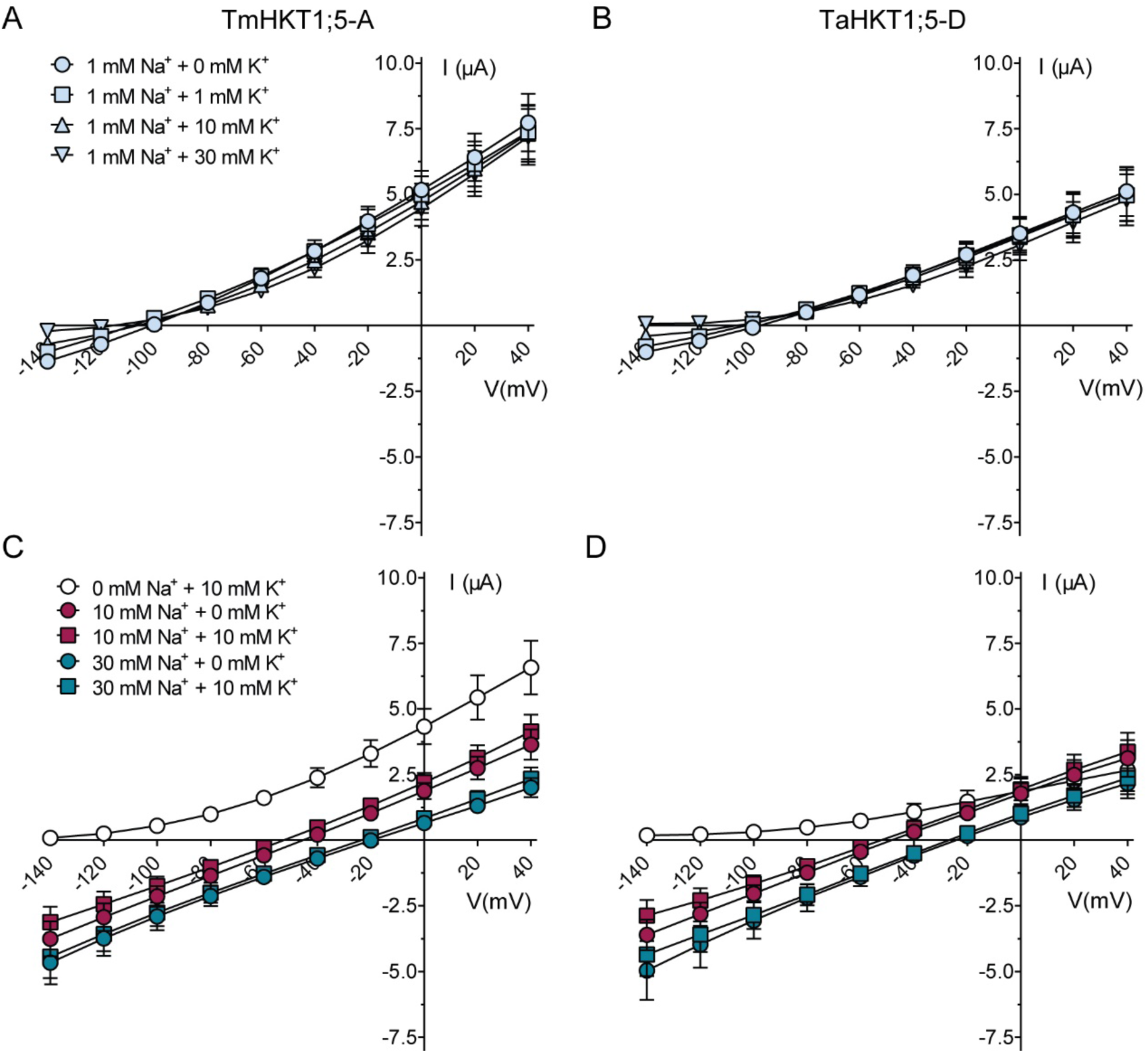
K^+^-sensitivity of Na^+^ currents carried by TmHKT1;5-A and TaHKT1;5-D. A-D, current-voltage (I/V) curves of Na^+^ currents recorded from *X. laevis* oocytes expressing *TmHKT1*;5-*A* (A, C) and *TaHKT1*;5-*D* (B, D) in 1 mM Na^+^ plus 0-30 mM K^+^ (A,B), or 0-30 mM Na^+^ solutions with or without 10 mM K^+^ (C,D). Mean ± S.E.M, n = 5-6.

**Figure 2.**
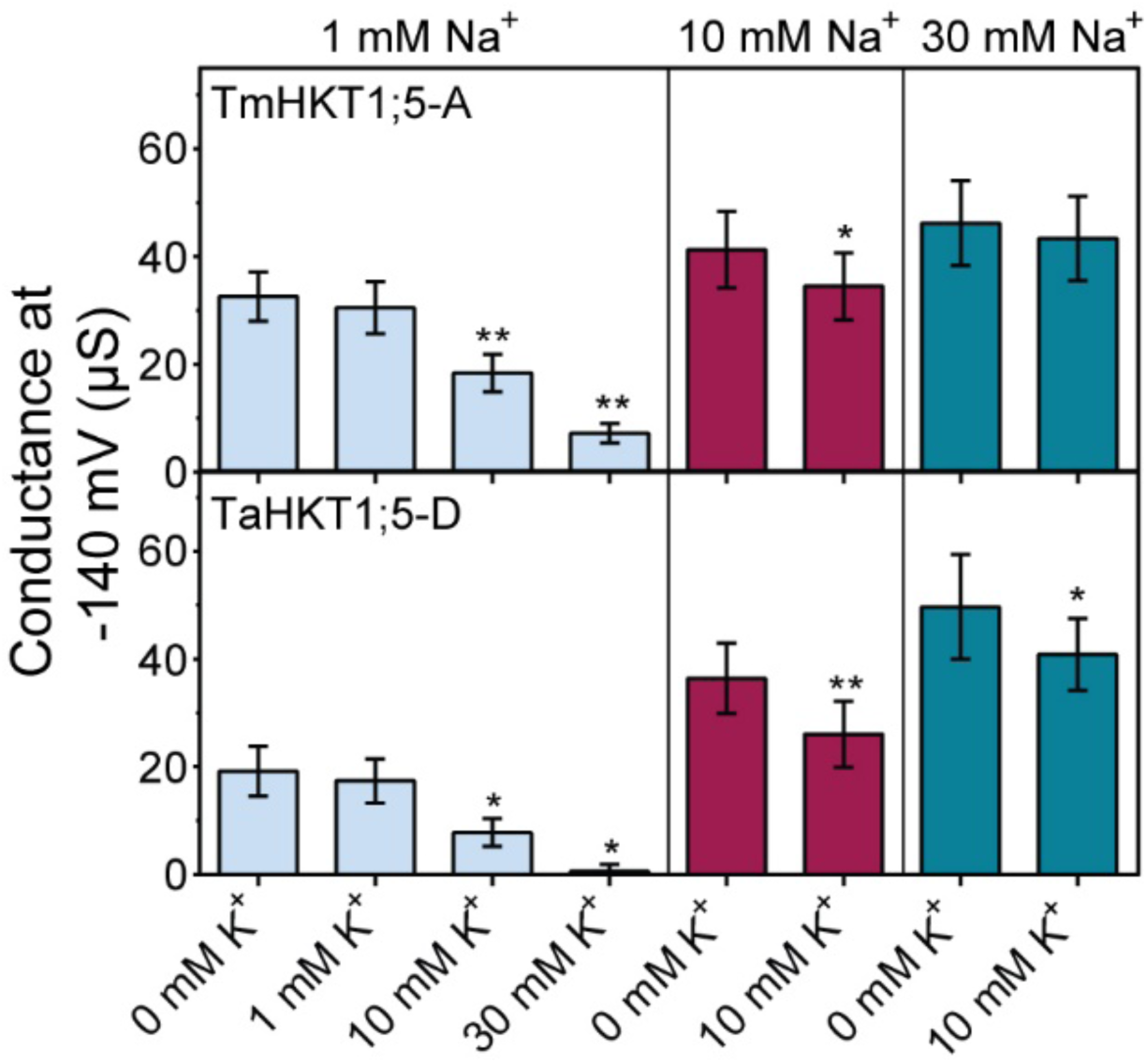
Channel conductance of TmHKT1;5-A and TaHKT1;5-D. The channel conductance of TmHKT1;5-A and TaHKT1;5-D was calculated from Figure 1, based on the slope of curve between −140 mV and −120 mV, statistical difference was determined by *Students’ t-test*, \**P < 0*.*05* and ** *P < 0*.*01*.

The increase in [K^+^]_ext_ from 0 to 10 mM did not significantly change the reversal potential of TmHKT1;5-A or TaHKT1;5-D; all reversal potentials were close to the theoretical equilibrium potential for Na^+^ under all conditions (Fig. 3A, 3B). Moreover, the observed shift in reversal potential were not significantly different from the theoretical Nernst shift for Na^+^ regardless of the presence or absence of 10 mM [K^+^]_ext_ (Fig. 3B). This suggests that neither of transporter proteins were permeable to Na^+^ and not to K^+^.

**Figure 3.**
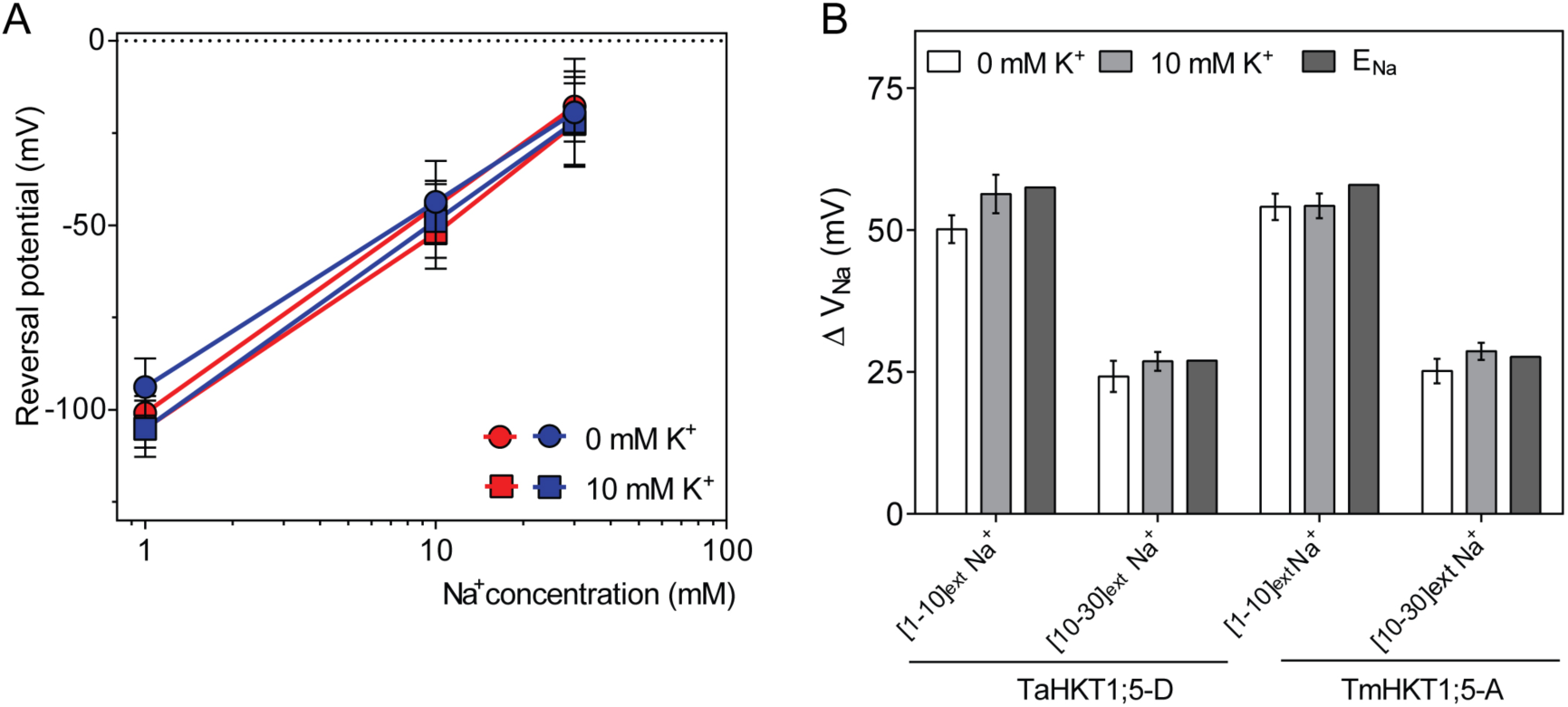
Shift in reversal potential of TmHKT1;5-A and TaHKT1;5-D in response to changes in [Na^+^]_ext_ and [K^+^]_ext_. A, Reversal potential of currents in *X. laevis* oocytes expressing *TmHKT1*;5-*A* (red) and *TaHKT1*;5-*D* (blue) derived from Figure. 1. B, Reversal potential of Na^+^ currents plotted with theoretical Nernst potential derived from internal concentrations measured in Munns et al. (2012) [19] and Byrt et al., (2014) [4]. Voltage at which current reversed in oocytes expressing *TmHKT1*;5-*A* and *TaHKT1*;5-*D* in solutions containing 1 mM to 10 mM Na^+^ ([1-10]_ext_ Na^+^) and 10 mM to 30 mM Na^+^ ([10-30]_ext_ Na^+^) with or without 10 mM [K^+^]_ext_, ΔV_Na_ of 0 mM or 10 mM K^+^ = experimentally measured shift in reversal potential for Na^+^ in 0 mM or 10 mM K^+^ conditions as indicated; ΔV_Na_ of E_Na_ = theoretical equilibrium (or Nernst) potential for Na^+^ was 57.7 mV and 26.9 mV, respectively for [1-10]_ext_ Na^+^ and [10-30]_ext_ Na^+^. One way ANOVA was performed for Figure 3B and no statistical differences were found between the theoretical and experimentally measured values.

### Dual affinity transport of Na^+^ by TmHTK1;5-A and TaHKT1;5-D

The affinity for Na^+^ transport of TmHKT1;5-A was previously reported to be ∼4-fold higher than the Na^+^-transport affinity of TaHKT1;5-D (4,6,19), which we confirm here (Fig. 4A, 4B). On closer examination, we found a property that has not been described before. When lower concentrations of [Na^+^]_ext_ were present it was clear that the concentration dependence of inward Na^+^ currents could be fitted by two components, for both TmHKT1;5-A and TaHKT1;5-D (Fig. 4). For TmHKT1;5-A, the high affinity component of Na^+^ transport had a K_*m*_ of 18.4 µM ± 0.1 µM when [Na^+^]_ext_ was below 0.1 mM (K_*m* (< 0.1 Na)_), whereas at concentrations of Na^+^ above 0.1 mM the K_*m*_ was 1.0 ± 0.2 mM (K_*m* (> 0.1 Na)_) (Fig. 4A). The K_*m*_ for the higher affinity component of inward Na^+^ transport for TaHKT1;5-D was 15.3 µM ± 0.9 µM when [Na^+^]_ext_ was lower than 0.1 mM (K_*m* (< 0.1 Na)_), whilst the lower affinity phase had a K_*m*_ of 4.3 ± 0.5 mM when [Na^+^]_ext_ was higher than 0.1 mM (K_*m* (> 0.1 Na)_) (Fig. 4B).

**Figure 4.**
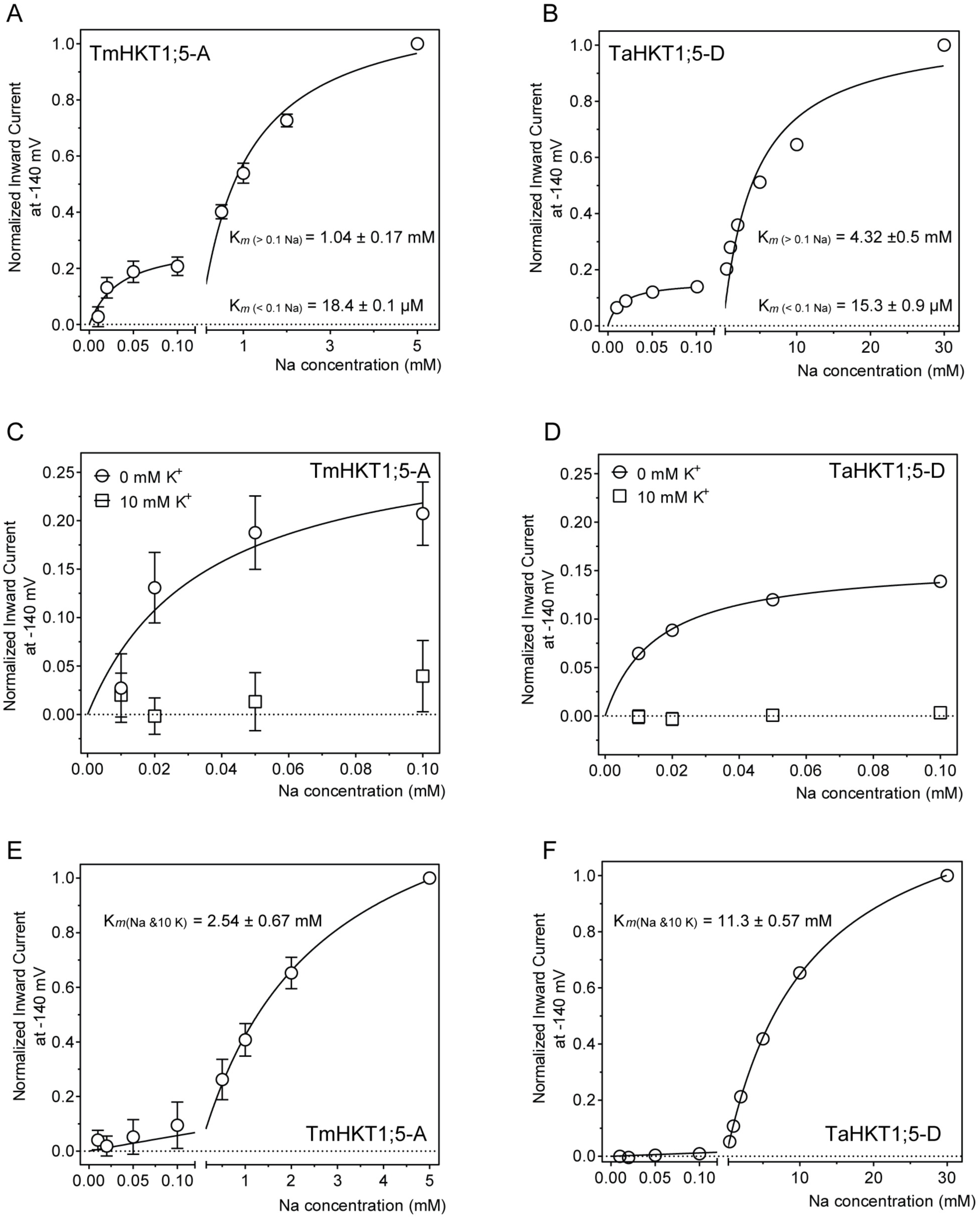
Transport affinity of TmHKT1;5-A and TaHKT1;5-D in *X. laevis* oocytes. A-B, dual Na^+^-transport affinity of TmHKT1;5-A and TaHKT1;5-D in *X. laevis* oocytes, plotted as normalised currents at −140 mV as shown against a series of Na^+^ glutamate solutions. C-D, high affinity phase of TmHKT1;5-A and TaHKT1;5-D in absence and presence of 10 mM [K^+^]_ext_. Data represented in A, B and C as mean ± S.E.M (n = 4 for A, n = 7 for B, n = 4 of 0 mM K^+^ and n = 5 of 10 mM K^+^ for C). E-F, Na^+^-transport affinity of TmHKT1;5-A and TaHKT1;5-D in *X. laevis* oocytes, plotted as normalised currents at −140 mV as shown against a series of Na^+^ glutamate solutions with an additional 10 mM K^+^ glutamate, n = 5 for E and n = 4 for F.

We examined the extent of K^+^-inhibition of both the lower and higher affinity components of the inward Na^+^ currents when [K^+^]_ext_ was 10 mM. The (normalised) inward Na^+^ current carried by both proteins was significantly suppressed by 10 mM [K^+^]_ext_, resulting in negligible inward Na^+^ transport by TmHKT1;5-A and TaHKT1;5-D in presence of [Na^+^]_ext_ below 0.1 mM (Fig. 4C and 4D). In the presence of 10 mM [K^+^]_ext_ it was now possible to fit the concentration dependence of inward current through TmHKT1;5-A and TaHKT1;5-D with one component. The K_*m*_ values of both TmHKT1;5-A and TaHKT1;5-D were decreased approximately by 2.5-fold by 10 mM [K^+^]_ext_ (Fig. 4E and 4F).

### Structural modelling of TaHKT1;5-D

Three-dimensional structural models of TaHKT1;5-D were generated to explore why K^+^ does not traverse the pore but instead blocks the Na^+^ currents, using the KtrB K^+^ transporter from *B. subtilis* as a template (6,24). The KtrB K^+^ protein was crystallised in the presence of KCl, hence the original structure contained a K^+^ ion in the selectivity filter region. This ion was substituted by Na^+^ during modelling of TaHKT1;5-D as it transports Na^+^ but not K^+^ (6). Ramachandran analysis indicated that the template and TaHKT1;5-D models generated in complex with Na^+^, and K^+^ had satisfactory stereo-chemical quality; Ramachandran plots showed only two residues positioned in disallowed regions (0.5% of all residues, except glycine and proline residues). Average G-factors (measures of correctness of dihedral angles and main-chain covalent forces of protein molecules) calculated by PROCHECK of the template and TaHKT1;5-D models with K^+^ and Na^+^ were 0.06. −0.18 and −0.13, respectively. The ProSa 2003 analysis of z-scores (−8.4; −6.0 and −5.8) indicated that template and modelled structures with K^+^ and Na^+^ had acceptable conformational energies.

The TaHKT1;5-D model in complex with Na^+^ showed that the Na^+^ ion coordination sphere had trigonal bipyramidal geometry, as defined by the Neighborhood algorithm (25), with coordination number 5. Na^+^ interacted with V76, S77, S78, N231, C232 and H351 (all carbonyl oxygens), that lies near the selectivity filter residues S78, G233, G353 and G457 (Fig. 5A). Notably, the carbonyl oxygen one of the selectivity filter residues (S78) directly participated in Na^+^ binding (Fig. 5A). Conversely, the TaHKT1;5-D model in complex with K^+^ showed that the K^+^ ion coordination sphere had octahedral geometry (25) with coordination number 6; K^+^ interacted with V76, N231, C232, H351, N455 and V456, that were also in the close neighbour of selectivity filter residues (Fig. 5B). However, in the second case, none of the selectivity filter residues participate in direct binding of K^+^. Ionic coordination spheres and Na^+^ and K^+^ distances of residues were evaluated using the Metal Binding Site Validation Server (25) that utilises more than 500 Protein Data Bank accession numbers. Ionic coordination spheres and binding modes were found to be acceptable, and ionic distances were within the following ranges: for Na^+^ ranged between 2.3 and 2.4 Å; for K^+^ were between 2.6 and 2.9 Å (Fig. 5, right panels).

**Figure 5.**
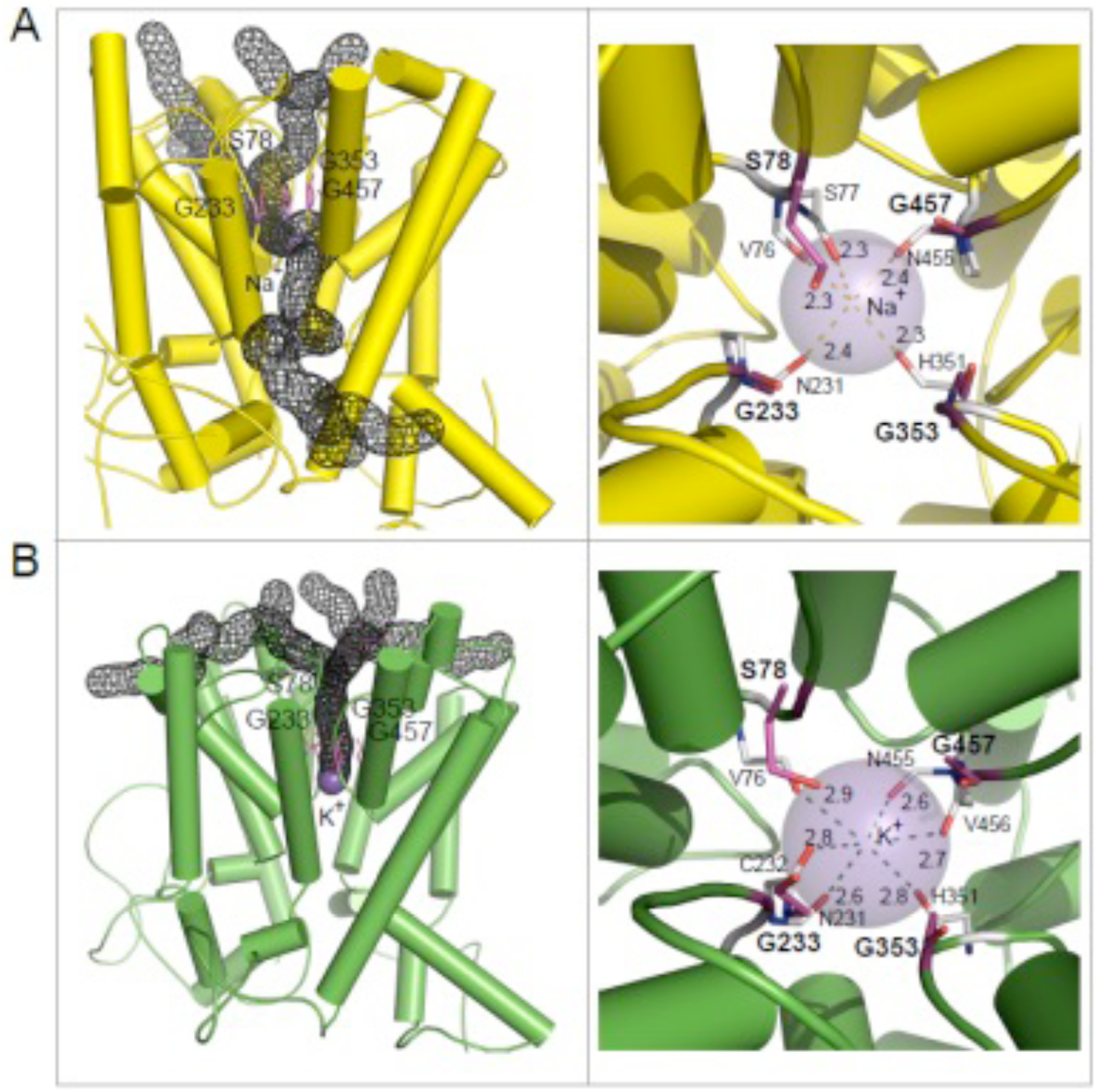
Molecular models of TaHKT1;5D in complex with Na^+^ (A) and K^+^ (B) ions. **(A-B; left panels)** Cartoon representations of TaHKT1;5-D illustrate overall protein folds with cylindrical α-helices, and translocating permeation channels (black mesh). Na^+^ (A) and K^+^ (B) ions are shown as purple spheres located within the selectivity filter residues S78, G233, G353 and G457 (coloured in cpk magenta), and other neighbouring residues. Na^+^ is likely to enter and exit the permeation trajectory from several points on both sides of the transporter, but always by-passes the selectivity filter constriction. K^+^ enters the selectivity filter and effectively blocks the permeation at the constriction bottleneck. For clarity, membrane α-helices are only shown. **(A-B; right panels)** Detailed views of coordination spheres of Na^+^ and K^+^ ions, and ionic interactions with residues adjoining the selectivity filter residues S78, G233, G353 and G457. The ionic distances of 2.3-2.8 Å for Na^+^ between V76, S77, S78, N231, C232 and H351, and of 2.6-2.9 Å for K^+^ between V76, N231, C232, H351, N455 and V456, are formed between carbonyl oxygens of residues (in sticks in atomic colours). The K^+^ ion effectively blocks the permeation channel through the protein pore at the constriction bottleneck point, contrary to Na^+^.

## Discussion

Oocytes expressing *TmHKT1;5-A* or *TaHKT1;5-D* have been previously reported to undergo no obvious change in their reversal potential regardless of the presence or absence of K^+^ in bath solution. Interestingly, both transporters had differential sensitivity to external [K^+^]_ext_. Whilst K^+^ block occurs for TmHKT1;5-A mediating inward transport it was not evident at [Na^+^]_ext_ above 10 mM, whereas this inhibition did happen for TaHKT1;5-D up to 30 mM [Na^+^]_ext_ (Fig. 2) (4,19). This differential inhibition effect of K^+^ upon Na^+^ transport at high [Na^+^]_ext_ by TaHKT1;5-D may result from a range of factors, that govern the rates of Na^+^ and K^+^ ion binding and subsequent Na^+^ transport. A variety of factors could control the transport of Na^+^ and the preliminary displacement of competing K^+^ with a consequent energetic cost, before Na^+^ passes desolvated or less-well hydrated than competing K^+^. Examples include free-energy variations between ions in binding sites relative to the corresponding quantity in bulk water, differences in selectivity filters solvent exposures (or dielectric constants), overall pore rigidity/stiffness (26), structural changes within funnels and selectivity filters due to residue re-orientation, differences in the hydration (coordination) numbers of Na^+^ and K^+^ (Na^+^ and K^+^ bind six and seven water molecules, respectively (27)) and the alterations on the rates of Na^+^ or K^+^ de-solvation (28). It is therefore plausible to expect that both ions would compete for binding sites within HKT funnel regions, yielding ion-protein binding sites with differential binding strengths.

Here, we conduct advanced 3D homology modelling and include explicit ionic radii of Na^+^ and K^+^ based on the CHARMM force field (29), rather than a traditional rigid body-type modelling for ligands. The aim has been to refine the previous structural model of HKT1;5 to show that the current model contains a pore that allows the passage of Na^+^ but not K^+^ (6,10,18). This is consistent with the functional observations in this and previous studies (4,19). Previously it was suggested that the Gly-Gly-Gly-Gly motif is important in coordinating the K^+^ ion in Ktr/TrK transporters and in conferring dual Na^+^-K^+^ transport in HKT2 transporters (16,30). Here our structural modelling suggests that the serine residue within the Ser-Gly-Gly-Gly selectivity motif directly interacts and binds with Na^+^ (Fig. 5B). Furthermore, our model suggests, despite ionic distances being shorter for Na^+^ than those for K^+^, that K^+^ could be bound more strongly due to a higher coordination pattern (Fig. 5A and 5B-right panels). Obviously, during binding, K^+^ takes an advantage of its larger ionic radius: 152 picometer (pm), compared to that of Na^+^ (116 pm); these parameters reflects empirical atomic radii of K (220 pm) and Na (180 pm). We assume that under the high concentrations of K^+^, this ion may outcompete Na^+^ and would become bound in the selectivity filter, thus effectively blocking Na^+^ transport. This is illustrated by calculating permeation channels in both complexes, using the Mole Voronoi algorithm, that predicts permeation trajectories and identifies path bottlenecks. From the calculated permeation paths, it could be deduced that the K^+^ ion effectively blocks the permeation channel through the major gating protein pore, contrary to Na^+^. It is also obvious that Na^+^ is likely to enter and exit the permeation trajectory from several points on both sides of the transporter, but always by-passes the selectivity filter constriction. Both ions arrive at the funnel of HKT proteins in solvated (hydrated) forms. While the hydration number for Na^+^ is 6, that for K^+^ is 7 at ambient temperature (27,31). The significance of the seven-fold water coordination for K^+^ is that this solvated complex may form a stronger hydrogen-bond interaction pattern (in funnels of HKT proteins), and would be de-solvated with a higher energy input and thus slower that the Na^+^ solvated complex (28). It is also plausible to expect that both solvated ion complexes would compete for binding sites within HKT funnel regions with different binding strengths. All in all, this would contribute to the blockage of the pore entry of HKT transporters by K^±^.

Potassium permeability for HKT1 transporters has only been reported once for the proteins that have been characterised in *X. laevis* oocytes. The transporters EcHKT1;1 and EcHKT1;2 allow permeation of both K^+^ and Na^+^ (15). Interestingly, when 3D models of the Na^+^ selective OsHKT1;5 and K^+^ permeable EcHKT1;2 were constructed and compared the pore size was predicted to be 0.2 Å larger in EcHKT1;2 (and that of OsHKT1;5 and TaHKT1;5-D), which would more than account for the weaker interactions with K^+^ and allow this larger ion to pass through the pore as well as Na^+^ (10). Without advanced 3D modelling of each and every transporter in the HKT family to show how the selectivity filter forms in the context of the remainder of the protein it would not be possible to predict which other HKT1s may allow passage of K^+^. Predictions based on sequence alone are insufficient to suggest how the two sets of proteins evolved their individual Na^+^ and K^+^ transport characteristics. An extension of the model to include other HKT proteins, a greater survey of structure-function relationships and mutations of key residues to predict functionally relevant residues and evolutionary relationships will be the focus of further studies.

Two affinity ranges for Na^+^ transport have been detected in the HKT family previously, but only for HKT2 transporters. OsHKT2;1, OsHKT2;2, OsHKT2;4 and HvHKT2;1(12,13,20,32-34). For instance, for OsHKT2;2, when K^+^ is present, the transport of Na^+^ is K^+^-dependent and displays a high affinity (K_*m*_ = 0.077 mM), whereas showing a lower affinity when absence of K^+^ (K_*m*_= 16 mM) (33,34). OsHKT2;1 also acquires two phases of Na^+^-transport affinity, whose K_*m*_ for Na^+^ is respectively 9.5 µM at [Na^+^]_ext_ below 1 mM and is 2.2 mM at [Na^+^]_ext_ > 1 mM (20). The K^+^ transport catalysed by HvHKT2;1 is affected by both [Na^+^]_ext_ and [K^+^]_ext_, such as a decrease in its K^+^ transport affinity from 30 µM down to 3.5 mM by [Na^+^]_ext_ increased from 0.5 mM to 30 mM and a greatly reduced channel conductivity for K^+^ by increasing [K^+^]_ext_ above ∼3 mM (35). This is the first time that dual affinity characteristics have been reported for the HKT1;*x* family (Fig. 4). Considering the relatively high homology between both clades (58% similarity/41% identity between TaHKT1;5 and TaHKT2;1) it is perhaps not surprising that both clades can confer dual affinity transport. How and why such ability has evolved and is conferred structurally is still to be determined.

In summary, we have detected that TaHKT1;5-D and TmHKT1;5-A can both conduct high and low affinity Na+ transport, both phases can be blocked by external K^+^ with the low affinity the low affinity component of Na^+^ transport for TmHKT1;5-A being less inhibited by external K^+^ than TaHKT1;5-D (Fig. 4). The two K_m_ at the high affinity phases were similar, but the K_m_ value for the low affinity component of TmHKT1;5-A was always ∼4 fold smaller regardless of the absence or presence of [K^+^]_ext_ (Fig. 4) (4,19). TmHKT1;5-A has been shown to be more effective in conferring shoot Na^+^ exclusion to wheat compared to TaHKT1;5-D (36). We propose that the higher affinity (with half-maximal activity nearer 1 mM compared to 4 mM), and a lower sensitivity to [K^+^]_ext_ are both characteristics that can help confer better shoot Na^+^ exclusion, which is a property that can lead to greater salt tolerance in the field for wheat (4,36,37).

## Materials and Methods

Brief methods for cloning of *TmHKT1;5-A* and *TaHKT1;5-D* as well as its functional characterisation in heterologous expression systems were described Munns et al. (4) and Byrt et al. (19). Further details are included here.

### Two-electrode voltage clamp recording in *X. laevis* oocytes

Oocyte recording followed the methods as described in Munns et al. (4) and Byrt et al. (19). Briefly, 46 nl/23 ng of cRNA or equal volumes of RNA-free water were injected into oocytes, followed by an incubation for 48 h before recording. Membrane currents were recorded in the HMg solution (6 mM MgCl_2_, 1.8 mM CaCl_2_, 10 mM MES and pH 6.5 adjusted with a TRIS base) ± Na^+^ glutamate and/or K^+^ glutamate as indicated in relevant Figures. All solution osmolarities were adjusted using mannitol at 220-240 mOsmol kg^−1^.

### Construction of 3D models of TaHKT1;5D in complex with Na^+^ and K^+^ ions

The most suitable template for TaHKT1;5D, the KtrB K^+^ transporter from *B. subtilis* (Protein Data Bank accession 4J7C, chain I) (24), was identified as previously described (6). The KtrB K^+^ protein was crystallised in the presence of KCl, hence the structure contains a K^+^ ion in the selectivity filter region. This ion was substituted by Na^+^ during modelling of TaHKT1;5D that transport Na^+^ at a greater rate than K^+^ (6). 3D structural models in complex with Na^+^, and K^+^ were generated using Modeller 9v16 (38) on a Linux station running the Fedora 12 operating system, as previously described (6,10,18). During modelling, attention was paid to ionic radii of Na^+^, and K^+^, whose topology parameters were taken from CHARMM (29). In each case, a total of 100 models were generated that were scored by Modeller using the modeller objective function (39), the discrete optimised protein energy function (40), PROCHECK (41), ProSa 2003 (42) and FoldX (43). Best scoring models constructed in Modeller 9v19 were further subjected to energy minimisation (knowledge-based Yasara2 forcefield with parameters for bond distances, planarity of peptide bonds, bond angles, Coulomb terms, dihedral angles, and van der Waals forces) (44), combined with the particle-mesh-Ewald (PME) energy function for long range electrostatics (cutoff 8.0 Å), to obtain smoothed electrostatic potentials. Structural images were generated with PyMOL Molecular Graphics System V1.8.2.0 (Schrődinger LLC, Portland, OR, USA). Permeation channels and path bottlenecks in both TaHKT1;5D ionic complexes were calculated using the Mole algorithm, embedded in PyMol (45), but using the x-y-z coordinates of Na^+^ or K^+^ as specifications for starting points.

